# Ultrasound elicits behavioral responses through mechanical effects on neurons and ion channels in a simple nervous system

**DOI:** 10.1101/104463

**Authors:** Jan Kubanek, Poojan Shukla, Alakananda Das, Stephen A. Baccus, Miriam B. Goodman

**Author notes:** **Corresponding author:** Miriam B. Goodman B-111 Beckman Center, 279 Campus Drive, Stanford, CA 94305.

## Abstract

Focused ultrasound has been shown to stimulate excitable cells, but the biophysical mechanisms behind this phenomenon remain poorly understood. To provide additional insight, we devised a behavioral-genetic assay applied to the well-characterized nervous system of *C. elegans* nematodes. We found that pulsed ultrasound elicits robust reversal behavior in wild-type animals in a pressure-, duration-, and pulse protocol-dependent manner. Responses were preserved in mutants unable to sense thermal fluctuations and absent in mutants lacking neurons required for mechanosensation. Additionally, we found that the worm‘s response to ultrasound pulses rests on the expression of MEC-4, a DEG/ENaC/ASIC ion channel required for touch sensation. Consistent with prior studies of MEC-4-dependent currents *in vivo*, the worm’s response was optimal for pulses repeated 300 to 1000 times per second. Based on these findings, we conclude that mechanical, rather than thermal stimulation accounts for behavioral responses. Further, we propose that acoustic radiation force governs the response to ultrasound in a manner that depends on the touch receptor neurons and MEC-4-dependent ion channels. Our findings illuminate a complete pathway of ultrasound action, from the forces generated by propagating ultrasound to an activation of a specific ion channel. The findings further highlight the importance of optimizing ultrasound pulsing protocols when stimulating neurons via ion channels with mechanosensitive properties.

**Significance Statement:** How ultrasound influences neurons and other excitable cells has remained a mystery for decades. Although it is widely understood that ultrasound can heat tissues and induce mechanical strain, whether or not neuronal activation depends on heat, mechanical force, or both physical factors is not known. We harnessed *C. elegans* nematodes and their extraordinary sensitivity to thermal and mechanical stimuli to address this question. Whereas thermosensory mutants respond to ultrasound similar to wild-type animals, mechanosensory mutants were insensitive to ultrasound stimulation. Additionally, stimulus parameters that accentuate mechanical effects were more effective than those producing more heat. These findings highlight a mechanical nature of the effect of ultrasound on neurons and suggest specific ways to optimize stimulation protocols in specific tissues.

## Introduction

Low-intensity, focused ultrasound affects the function of excitable cells in the central (Fry and others, 1958; Meyers et al., 1959; Foster and Wiederhold, 1978; Gavrilov et al., 1996; Tufail et al., 2011; Yoo et al., 2011; Deffieux et al., 2013; Menz et al., 2013; King et al., 2013) and peripheral (Mihran et al., 1990; Tsui et al., 2005; Colucci et al., 2009) nervous systems, and the heart (Harvey, 1929; Buiochi et al., 2012). Because it propagates deep into tissue while retaining spatial focus, it has garnered considerable attention for its potential as a non-invasive tool for stimulation of the brain and the heart (Tufail et al., 2011; Lee et al., 2016). Despite these investigations, how ultrasound stimulates excitable cells has been a mystery since the discovery of its stimulatory effects more than eight decades ago (Harvey, 1929).

Physical mechanisms for ultrasound-dependent tissue excitation have been divided into thermal and non-thermal effects (Dalecki, 2004; Sassaroli and Vykhodtseva, 2016; Naor et al., 2016; Ye et al., 2016). The latter category encompasses mechanical effects such as membrane oscillation, cavitation, or radiation force. Ultrasound stimulation generally occurs under conditions that heat tissues by less than 1°C (Tufail et al., 2010; Yoo et al., 2011; Menz et al., 2013). Nonetheless, even small temperature changes could activate certain classes of ion channels, such as temperature-sensitive TRP cation channels (Diaz-Franulic et al., 2016) and TREK potassium channels (Schneider et al., 2014). Moreover, the rate of temperature change may also contribute to the stimulatory effects (Rabbitt et al., 2016).

The mechanical effects of ultrasound could be converted into ionic currents and changes in electrical excitability by increasing mechanical strain in a manner that directly activates ion channels (Tyler, 2011) or by changes in membrane thickness and capacitance (Plaksin et al., 2014). For instance, increases in membrane tension are thought to catalyze conformational change and activate ion channels in which the open state has a larger cross-sectional area than the closed state (Sukharev and Corey, 2004). Thus, if ultrasound stimulation were to change membrane tension, then it could alter the activity of mechanosensitive channels including voltage-gated sodium and potassium channels that exhibit membrane tension-dependent gating (Beyder et al., 2010; Brohawn et al., 2014). Consistent with this idea, ultrasound stimulation has been shown to increase currents carried by two-pore domain (K2P) potassium channels and voltage-gated sodium channels expressed in Xenopus oocytes (Kubanek et al., 2016).

This body of work has raised the question whether or not ultrasound affects nervous systems via thermal or non-thermal effects (Iversen et al., 2017) and, if non-thermal effects are dominant, how is ultrasound transduced into membrane strain (tension) and channel activation. To contribute additional insight into these questions, we developed a behavioral-genetic assay using *C. elegans* roundworms. The nervous system of this animal has been characterized in its entirety, the animal performs simple behaviors that are easily monitored using video tracking and machine vision tools (Husson et al., 2005), and the model is amenable to experimental studies involving tens to hundreds of individual animals. Two additional aspects of the worm’s biology are exploited in this study. First, *C. elegans* has extraordinarily sensitive thermosensory and mechanosensory neurons able to detect thermal fluctuations of 0.05°C or less (Ramot et al., 2008) and applied forces of 50 nN (O’Hagan et al., 2005), respectively. Second, mutants exist in which thermosensory and mechanosensory neurons are disabled independently.

We found that pulsed ultrasound stimulation evokes avoidance responses whose probability increases with acoustic pressure and stimulus duration and shows optima for specific ultrasound pulsing protocols. Mutants lacking neurons required for thermosensation responded similar to wild type animals, whereas those lacking neurons or ion channels required for mechanosensation failed to mount avoidance responses to ultrasound stimulation. The mechanical nature of the effect led us to a detailed characterization of the involved neurons and ion channels as well as a characterization of the optimal stimulation parameters.

## Materials and Methods

### Animals and strains

The following strains were used in this study: *N2(RRID: WB - STRAIN: N2(ancestral)*);CB1338 *mec-3(e1338)* IV (RRID:WB-STRAIN:CB1338); CB1611 *mec-4(e1611)* X (RRID:WB-STRAIN:CB1611); TU253 *mec-4(u253)* X (RRID:WB-STRAIN:TU253); IK597 *gcy-23(nj37)gcy-8(oy44)gcy-18(nj38)*) IV (RRID:WB-STRAIN:IK597); VC1141 *trp-4(ok1605)* I (RRID:WB-STRAIN:VC1141); VC818 *trp-4(gk341)* I (RRID:WB-STRAIN:VC818); TQ296 *trp-4(sy695)* I (RRID:WB-STRAIN:TQ296); GN716 *trp-4(ok1605)* I, outcrossed four times from VC1141. All mutants were derived from the N2 (Bristol) background, which serves as the wild-type strain in this study. Except for GN716, which was prepared explicitly for this study, and TQ296, which was a gift of X. Z. S. Xu (U Michigan), strains were obtained either from a frozen repository maintained in the Goodman lab or from the Caenorhabditis Genetics Center.

The three *trp-4* alleles we studied all encode deletion or null alleles of the *trp-4* gene, which encodes the key pore-forming subunit of a mechanosensory ion channel expressed in the CEP texture-sensing neurons (Kang et al., 2010). TRP-4 is orthologous to the mechanically-gated NOMPC channel from Drosophila.

The *ok1605* allele encodes an in-frame, 1-kb deletion that removes exons 12-14 of the *trp-4* gene and is predicted to result in the loss of ankyrin repeats 16-21 from the TRP-4 protein. The *gk341* allele contains a small deletion encompassing exon 2 and is predicted to cause a frame-shift in the transcript and the introduction of an early stop codon. The *sy695* allele contains an unmapped 3kb deletion in the 3’ region of the gene. This deletion is thought to disrupt the transmembrane pore-forming domain of TRP-4 (Li et al., 2006). The GN716 *trp-4(ok1605)* strain was derived by out-crossing VC1141 *trp-4(ok1605)* with wild type (N2) in four rounds. We used PCR to verify that all *trp-4* mutant strains harbored the expected genetic deletions in the *trp-4* locus.

### Imaging and transducer control

For each assay, we transferred a single adult animal from a growth plate to a 4 mm-thick NGM agar slab that was free of bacteria. Because the agar slab consists of a 2% agar solution in saline, it is a good approximation of the acoustic properties of many biological tissues (Altman et al., 1974). To retain animals within the camera’s field of view, we created a boundary consisting of a filter paper ring saturated by a copper sulfate (500 mM) solution, as described (Ramot et al., 2008).

To deliver ultrasound stimuli, we used a commercially-available piezoelectric ultrasonic transducer (A327S-SU-CF1.00IN-PTF, Olympus, 1-inch line-focused) positioned 1 inch (2.54 cm) below the top of the agar slab and oriented perpendicular to the surface of the agar slab (Fig. 1A). We filled the space between the transducer surface and the bottom of the agar slab with degassed water, contained within a plastic cone mounted on the transducer. The water was degassed by boiling for 30 min and stored in air-tight tubes. The slab did not appear to attenuate ultrasound pressure, according to measurements with a hydrophone (not shown).

We used oblique illumination via a circular array (20 cm in diameter) of red LEDs to provide the optical contrast between animals and the surface of the agar slab needed to track animal movement using the Parallel Worm Tracker (Ramot et al., 2008). The image was magnified 3x (zoom lens, Navitar). The camera’s chip field of view was 5.6 × 4.2 mm. Image contrast was optimal when the plane of the LED array was about 1 cm above the top of the agar slab. We also used a blue LED, controlled by an Arduino Uno board and mounted 5 cm above the agar slab, to deliver an optical signal synchronized to the stimulus onset.

To generate signals driving the ultrasound transducer, we used a function generator (HP 8116A, Hewlett-Packard, Palo Alto, CA) controlled by an Arduino board (Uno) and an amplifier with a gain of 50 dB (ENI-240L, ENI, Rochester, NY). The acoustic pressures we generated were measured in free field using a calibrated hydrophone (HGL-0200, Onda, Sunnyvale, CA) combined with a pre-amplifier (AG-2020, Onda). The hydrophone measurements were performed at the peak spatial pressure. The hydrophone manufacturer’s calibration values around the frequency of 10 MHz were steady and showed only minimal level of noise.

### Behavioral recordings

For each trial, a freely moving animal was monitored via a digital video camera (SME-B050-U, Mightex) operated in a live-video mode until it approached the ultrasound focus head first. We started recording videos approximately 5 s before the predicted approach to the ultrasound focus and continued recording for roughly 10 s following the delivery of each stimulus. Each individual animal was tested in 10 trials with an inter-trial interval of at least 20 s. All animals were hermaphrodites, assayed blind to genotype and as adults. We applied stimuli at the following pressures by controlling the output of the function generator (voltage in mV, pressure in MPa): 0 (*aka* sham treatment, the amplifier was operated but not connected to the transducer), 0.2 (60), 0.4 (120), 0.6 (180), 0.8 (240), and 1.0 (300) MPa. We were not able to deliver stimuli more intense than 1.0 MPa because the transducer was damaged by sustained operation at this level. The protocol for determining the effects of stimulus duration, duty cycle, and pulse repetition frequency was analogous except that we varied the levels of the respective quantities.

Each animal’s movement was recorded at 20 frames per second at a resolution of 576 × 592 pixels. We recorded roughly 350 frames per video. The resolution and frame-rate were chosen to be high enough to provide reliable movement characterization while maintaining acceptable size of the stored videos.

### Quantification of response frequency and baseline response frequency

To detect *bona fide* ultrasound-evoked reversals, we measured the velocity vector over the interval from 250 ms to 1 s following the ultrasound onset and that during a 1 s period immediately preceding the ultrasound onset. Next, we computed the vector difference, and evaluated the magnitude of that difference. We asked whether this metric differed from the null distribution constructed over all baseline recordings (same time windows, just shifted 1 s back in time so that there could be no effect of ultrasound) available for a given animal. If the vector difference was unlikely (*p* < 0.01) to have been drawn from the null distribution, we classified the response as a reversal. We computed the proportion of significant responses over the 10 stimulus repetitions for each animal and refer to this metric as the response frequency.

The computation of the baseline response frequency (dashed lines in the plots) was analogous to the computation of the response frequency with the exception that the metrics were taken in time windows before the ultrasound could have any impact (i.e., before the ultrasound was turned on). In particular, the velocity difference was computed by comparing a 1 s time window immediately preceding the ultrasound to a 1 s time window preceding the ultrasound onset by 1 s. The baseline distribution used the same time windows, just shifted back in time by 1 s. The baseline response frequency was indistinguishable across the tested animal strains (*F*_4,95_ = 0.28, *p* = 0.90, one-way ANOVA).

### Simulation of the relationship between reversal frequency and duty cycle

The simulation shown in Fig. 5B was derived as follows. First, we assumed that the signal relevant to modulation of reversal behavior is the envelope modulating the carrier frequency of the ultrasound transducer. Next, for each duty cycle value we tested, we converted this signal into the frequency domain using the function `fft` in Matlab. Finally, this signal was convolved with the filter computed by Eastwood et al. (Eastwood et al., 2015) and the effective (rms) value of the resulting signal was taken as the model’s output. Thus, this simulation rests on the delivered stimulus waveform and the previously proposed mechanical filter. The only adjustment was a linear scaling factor used to plot the results on common graph. We note that filter developed by (Eastwood et al., 2015) was defined over the range from 1 Hz to 3 kHz, which we extrapolated to 10 kHz.

### Temperature measurements

We used a dual sensing fiber-optic hydrophone (Precision Acoustics, Dorchester, UK) to measure temperature (Morris et al., 2009). This device uses optical interferometry to record acoustically- and thermally-induced changes in thickness of its sensing membrane by ultrasound. When performing the measurements, we immersed the tip of the optic fibre into the agar at the location of maximal pressure. The device, which recorded the temperature at a 200 Hz sampling rate, rapidly registered the changes in temperature to ultrasound onset. As expected, temperatures were highest at the end of the ultrasound burst; consequently, we report the difference in temperature between the start and end of the ultrasound burst.

### Simulation of acoustic radiation force

We used the k-Wave simulation tool (Treeby and Cox, 2010) to estimate the acoustic radiation force that acted on the animals in our setup. The simulation used the same geometry and media as our setup, including water, agar (4 mm thick), worm on the agar (0.05 × 0.05 × 1.0 mm), and air. The speed of sound (m s^-1^), density (kg m^-3^), and the acoustic attenuation coefficient (dB cm^-1^ MHz^-1^) for these media were respectively set to: (1540,1000, 0.0022), (1548,1000, 0.40), (1562,1081,1.2), (343,1.2,1.6) (Barber et al., 1970; Kremkau et al., 1981). The simulation grid was computed in steps of 1 ns in time and 10 *µ*m in space. We set the stimulus level such that the simulated pressure field had an amplitude of 1 MPa at focus. We obtained the steady-state time-average radiation force density field from the steady-state time-average intensity field provided by the simulation. To obtain the net radiation force, we integrated the force density field over a volume that approximated the animal‘s head (0.05 × 0.05 × 0.2 mm). This resulted in a net force magnitude of 873 nN. Integrating the force density over a larger volume had only a small influence on the results since the force field was strongest near the animal’s head (Fig. 1A). In Fig. 1B, the force density is integrated within cubes of 0.05 × 0.05 × 0.05 mm, using acoustic parameters of the worm.

### Experimental Design and Statistical Analysis

The *C. elegans* hermaphrodites used in this study were age-synchronized by hypochlorite treatment (Stiernagle, 2006) and cultivated at 15 or 20°C. There were no detectable effects of cultivation temperature, ambient temperature, or humidity on the ultrasound-evoked responses, which were collected over a period of months by two members of the research team. All behavioral recordings were performed blind to genotype and we determined whether or not a given trial included a reversal event as described above. The response rate was computed as the fraction of ten trials that evoked a reversal and results were pooled from *n* = 20 *C. elegans* subjects.

We used ANOVAs to assess the effect of stimulus pressure, duration, pulse repetition frequency, and duty cycle. In this linear model (Fig. 3, Fig. 5), the dependent variable is the response frequency as described above. The independent variable (the one factor) is pressure (Fig. 3), or pulse repetition frequency or duty cycle (Fig. 5). We report the *p*-values as well as the *F*-statistic and the degrees of freedom of the tests.

## Results

As a first step toward determining how animals and excitable tissues detect and respond to ultrasound stimulation, we placed single adult wild-type (N2) nematodes on sterile agar slabs and tracked their movement using a digital video camera and the Parallel Worm Tracker (Ramot et al., 2008), adapted for tracking single animals (Fig. 1A). We subjected each animal to pulsed ultrasound (10 MHz frequency, 200 ms duration, 1 kHz pulse repetition frequency at 50% duty cycle, Fig. 1B) when it approached the ultrasound focus and found that this stimulus elicits similar reversal behaviors over the course of ten trials (Fig. 2). The effect was due to ultrasound stimulation *per se* since sham stimuli (0 MPa) did not increase reversals above their basal or unstimulated rate (Fig. 2A, B).

**Figure 1.**
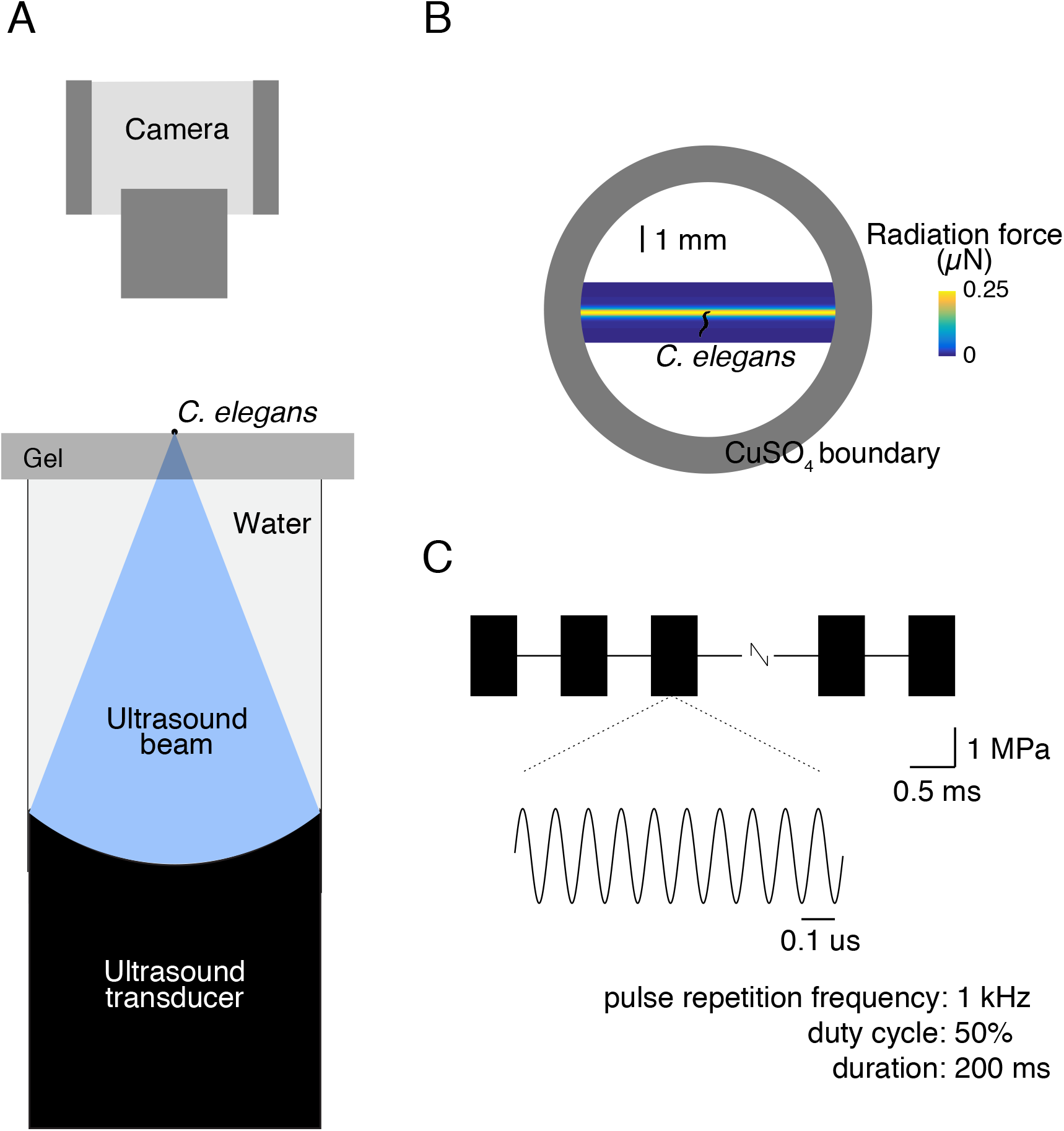
A system for delivering pulsed ultrasound to *C. elegans* nematodes. (A) Schematic side view of the set-up, showing a single wild-type adult hermaphrodite crawling on the surface of agar slab, tracked by a digital video camera, and maintained within the field of view by a copper sulfate boundary. A piezoelectric ultrasound transducer (10 MHz carrier frequency, line-focused) is coupled directly to the bottom of the agar slab by a column of degassed water. (B) Simulation of the distribution of radiation force expected from the line-focused 10MHz ultrasound transducer (see Methods). Animals were stimulated only during forward movement, as they entered the zone corresponding to the highest expected pressure. (C) Schematic of a typical stimulus consisting of ultrasound pulses delivered at pulse repetition frequency of 1 kHz for a total duration of 200ms at 50% duty cycle. In this study, we systematically varied the applied pressure, pulse duration, pulse repetition frequency, and duty cycle.

**Figure 2.**
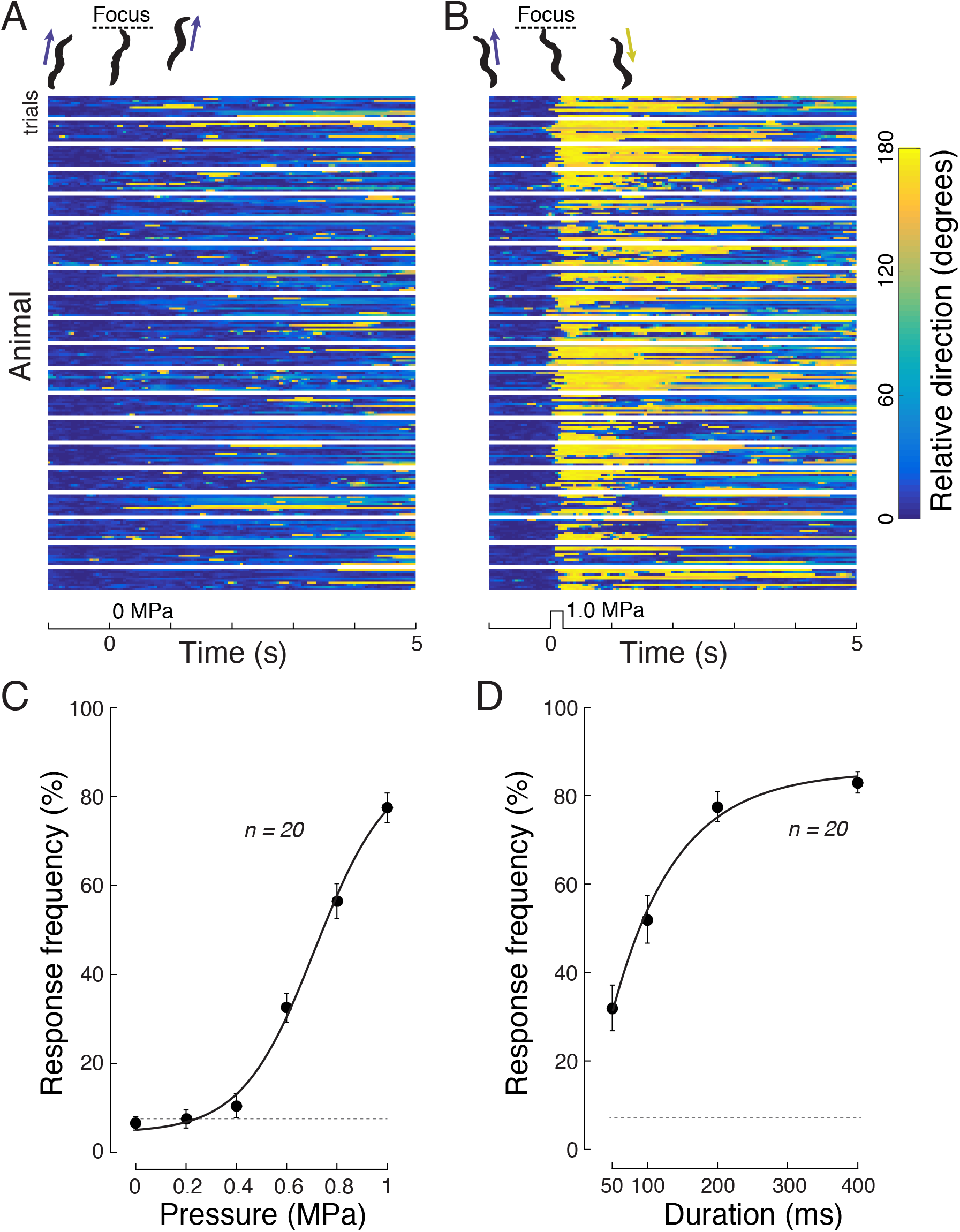
Ultrasound elicits reversal behavior in a pressure and stimulus time-dependent manner in wild-type *C. elegans*. (A, B) Raster plots showing the response of 20 animals (10 trials/animals) to a 200-ms sham stimulus (0 MPa pressure, Panel A) and a bona fide stimulus (1.0 MPa, Panel B). Heading angle is encoded in color such that headings similar to the average angle in the 1 s window immediately preceding stimulus onset are blue and reversals are encoded in yellow. Rows correspond to single trials and blocks are ten trials delivered to each animal; traces were smoothed with a zero-lag rectangular sliding 150-ms window. The silhouettes (top) depict representative responses to sham (A) and 1.0 MPa stimuli (B). (C) Reversal frequency increases with applied pressure. Points are mean±s.e.m. (*n* = 20) for animals stimulated at each of the six pressure values for a total of 10 trials. The solid line is a Boltzmann fit to the data with an *P*_1/2_ of 0.71 MPa, a slope factor of 0.15, and a maximum probability of 83%. The dotted line is the unstimulated reversal rate. Stimulus parameters: 1kHz, 50% duty cycle, 200ms pulse duration, variable pressure. (D) Reversal probability increases with stimulus duration. Points are means.e.m. (*n*=20) and the smooth line is an exponential fit to the data with a time constant of 90 ms. Stimulus parameters: 1 kHz, 50% duty cycle, variable pulse duration, 1.0 MPa pressure. In both C and D, the dotted line represents baseline rate of responding (see Materials and Methods). Smooth line is an exponential fit to the data with a time constant of 90 ms.

Behavioral responses were robust across trials for a given individual and among all animals tested (Fig. 2B). We determined whether each animal’s response to ultrasound stimulation was statistically different from spontaneous changes in direction (Croll, 1975) (see Materials and Methods for details). For each animal, we quantified the proportion of significant responses over the 10 stimulus trials, and refer to this metric as response frequency.

The response frequency increased with increasing ultrasound pressure (Fig. 2C) and was indistinguishable from the spontaneous reversal rate for the sham stimulus (Fig. 2A, dotted line; *p* = 0.52, t-test, *n* = 20). The reversal rate increased above the baseline at a pressure of 0.6 MPa (*p* < 10^−6^). At 1 MPa (Fig. 2B) there was a significant response in 77.5% of trials, on average. The relationship between response frequency and applied pressure was well-described by a sigmoid function: 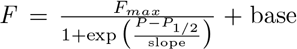, where *F* is the response frequency, *P* is the pressure, and *base* is the spontaneous reversal frequency. For wild-type animals stimulated by a 200-ms pulse (1kHz pulse repetition frequency, 50% duty cycle), the best fit parameters were *F*_max_ = 83%; *P*_1/2_ = 0.71 MPa, slope = 0.15, base = 5%. Thus, the half-activation pressure equals 0.71 MPa. A one-way ANOVA also detected a significant modulation of the response frequency by pressure (*F*_5,114_ = 103.4, *p* < 10^−39^), reinforcing the idea that the probability of ultrasound-induced reversal depends on stimulus pressure.

We also tested the effect of varying the total duration of the ultrasound stimulus (Fig. 2D), holding all other parameters (i.e. pressure, duty cycle, pulse repetition frequency) constant. In agreement with a previous study (Ibsen et al., 2015), responses were weak or absent when the stimulus was brief. There was a significant modulation of the response frequency by stimulus duration (one-way ANOVA, *F*_3,76_ = 30.8, *p* < 10^−12^). Stimuli of 100 ms in duration or longer produced substantial effects. The response frequency did not increase substantially beyond stimulus duration of 200 ms (response frequency at 200 ms versus 400 ms: *p* = 0.24, paired t-test, *n* = 20). Therefore, we used a stimulus duration of 200 ms for subsequent experiments.

In principle, ultrasound-evoked behaviors could depend on thermosensation, mechanosensation, or both. We used a genetic approach to distinguish among these possibilities, leveraging mutants deficient in thermosensation or mechanosensation. To test for thermal effects, we compared ultrasound-evoked behaviors in wild-type and *gcy-23(nj37)gcy-8(oy44)gcy-18(nj38)* that lack a trio of receptor guanylate cyclases expressed exclusively in the AFD thermoreceptor neurons and are defective in thermotaxis (Garrity et al., 2010; Glauser and Goodman, 2016). Although these mutants have an intact AFD thermoreceptor neuron, they lack the ability to sense tiny (<0.05°C) thermal fluctuations in temperature (Ramot et al., 2008; Wasserman et al., 2011). As shown in Fig. 3A, the response of these thermosensory-defective mutants was indistinguishable from that of wild type animals. The mutants retained modulation by stimulus pressure, as assessed by one-way ANOVA (*F*_5,114_ = 80.7, *p* < 10^−35^). Furthermore, as expected from the plot, a two-way ANOVA with factors animal strain and pressure failed to detect a significant difference between the strains (*F*_1,228_ = 0.02, *p* = 0.89) as well as the strain × pressure interaction (*F*_5,228_ = 1.40, *p* = 0.23). Thus, the ability to sense tiny (<0.05°C) thermal fluctuations is not required for ultrasound-induced reversal behaviors.

Having established that thermosensation is dispensable for ultrasound-evoked reversals, we compared responses in wild-type animals and mutants defective in mechanosensation. Specifically, we sought to quantify ultrasound-evoked responses in mutants in which selected mechanoreceptor neurons fail to properly differentiate during development, degenerate, or lack essential, pore-forming subunits of known sensory mechano-electrical transduction (MeT) channels: MEC-4 (O’Hagan et al., 2005), TRP-4 (Kang et al., 2010). The goal of these experiments was to identify the neurons most likely to serve as the first responders to ultrasound stimulation and to determine whether or not such sensitivity relied upon known MeT channels.

The *mec-3(e1338)* fail to generate three sets of neurons known to participate in gentle and harsh touch sensation (TRN, PVD, FLP) mutants and are insensitive to both gentle and harsh touch (Way and Chalfie, 1989). The six touch receptor neurons (TRNs: ALML/R, AVM, PLML/R, PVM) are required for sensing gentle touch and the two pairs of multidendritic PVD and FLP neurons act as polymodal nociceptors (Schafer, 2015). We found that *mec-3* mutants are insensitive to ultrasound stimulation (Fig. 3B): the *mec-3* mutants showed no significant modulation of the response frequency by pressure (*F*_5,114_ = 1.18, *p* = 0.32, one-way ANOVA). A two-way ANOVA detected both a highly significant difference between the strains (*F*_1,228_ = 246.1, *p* < 10^−37^) and a highly significant strain × pressure interaction (*F*_5,228_ = 56.8, *p* < 10^−37^). This effect was specific to ultrasound-evoked reversals; mutants moved at an average speed that was indistinguishable from wild-type animals (speed measured during 1 s period preceding ultrasound onset; wildtype: 0.21 mm/s; *mec-3*: 0.17 mm/s; *p* = 0.11, *n* = 20, t-test). These average speeds are within the range of values reported previously for wild-type animals (Ramot et al., 2008). This result shows that the *mec-3*-dependent mechanoreceptor neurons are required for ultrasound-evoked reversals and suggests that ultrasound can exert forces on neural tissue sufficient to activate these neurons.

We narrowed the search to a subset of the *mec-3*-dependent mechanoreceptor neurons by testing ultrasound-evoked behavior in *mec-4(e1611)* mutants in which the six TRNs degenerate and the PVD and FLP mechanoreceptor neurons are intact (Driscoll and Chalfie, 1991). As found in *mec-3* mutants, ultrasound failed to evoked reversals in *mec-4(e1611)* (Fig. 3C) and there was no significant modulation of the response frequency by the ultrasound pressure amplitude in these animals (Fig. 3C; *F*_5,114_ = 1.47, *p* = 0.20). Moreover, a two-way ANOVA detected a significant difference between the mutant and wild-type strains and a highly significant strain × pressure interaction (both *p* < 10^−36^). Thus, the TRN neurons, which can detect forces as small as 50 nN (O’Hagan et al., 2005), are required for behavioral responses to ultrasound stimulation in *C. elegans*.

**Figure 3.**
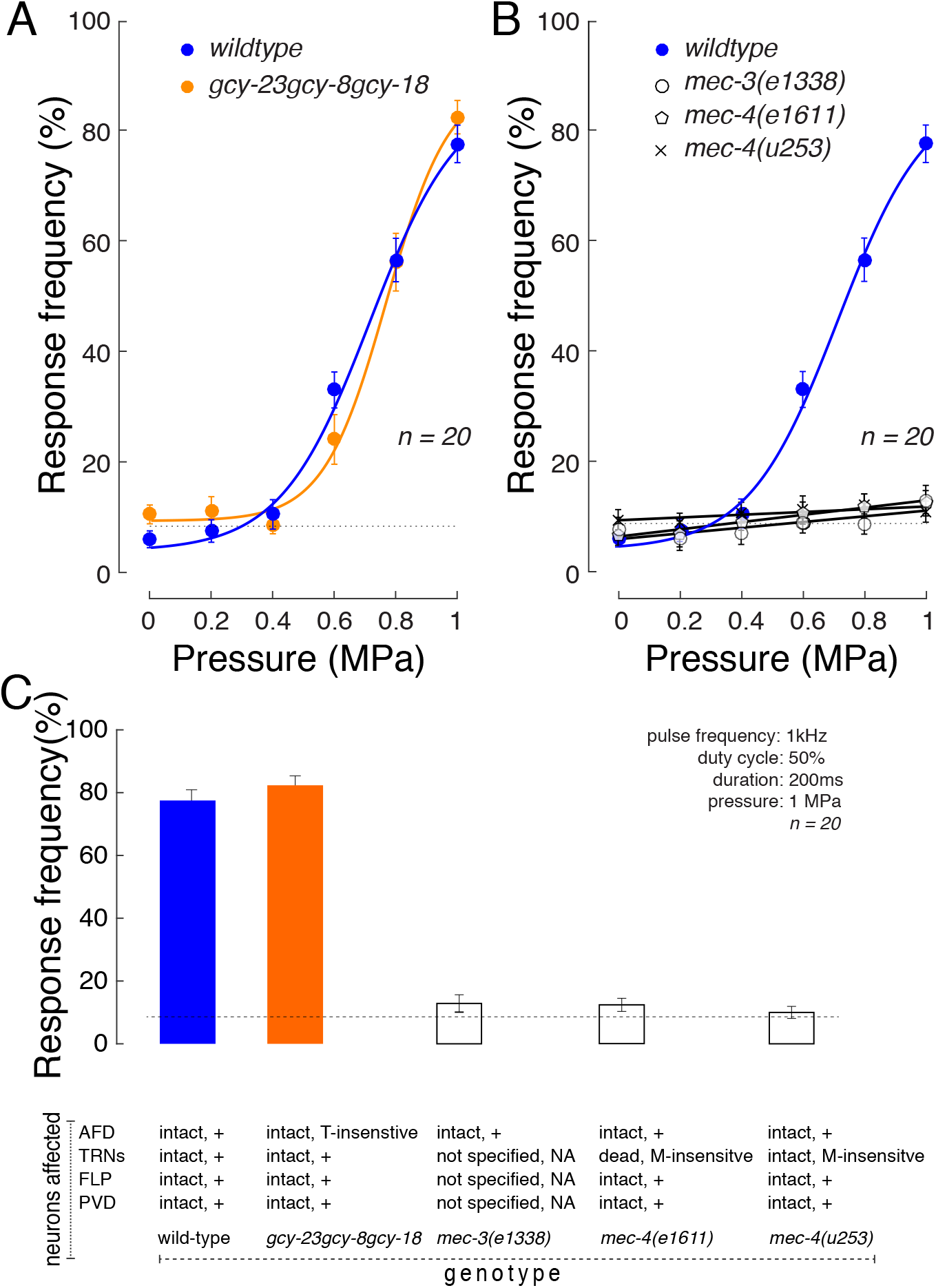
Loss of mechanosensation, but not thermosensation disrupts ultrasound-evoked reversals. (A, B) Pressure-response curves of wild-type N2 animals (blue) compared to a thermosensation-defective mutant (orange) and three mechanosensation-defective mutants (black), *mec-3(e1338), mec-4(e1611)*, and *mec-4(u253)*. Points are mean ±s.e.m. (*n* = 20 animals tested in 10 trials/animals) and smooth lines are fit to the data according to a sigmoidal function. The data and fit for wild-type are the same as in Fig. 2C. Fitting parameters for *gcy-23(nj37)gcy-8(oy44)gcy-18(nj38)* are (*F*_max_, *P*_1/2_, slope, base): 80%, 0.76 MPa, 0.10, 9%. Dotted lines are the average baseline response rate (see Materials and Methods) for each case; there was no significant effect of genotype on baseline reversal rates. Stimulus parameters: pulse frequency: 1kHz; duty cycle: 50%; duration: 200ms; pressure: variable. (C) Response rate as a function of genotype. Bars are the mean (s.e.m., *n*=20) reversal rate. Annotations below the graph indicate the nature of the known sensory deficit associated with each genotype (see Text for detail). Animals tested as young adult hermaphrodites and blind to genotype.

Next, we investigated proteins expressed in the TRN neurons that might mediate the effect. Of particular interest, the TRNs express *mec-4* which encodes a nonvoltage-gated sodium channel of the DEG/ENaC/ASIC family required for touch-evoked reversals. MEC-4 is expressed exclusively in the TRNs and is an essential pore-forming subunit of the mechanosensitive ion channel activated by mechanical loads applied directly to the animal‘s skin (O’Hagan et al., 2005). Like *mec-3* and *mec-4(e1611)* mutants, *mec-4(u253)* null mutants are insensitive to ultrasound stimulation (Fig. 3D). These animals have intact TRNs, but lack the MEC-4 protein required for mechanotransduction and showed no significant modulation of the response frequency by the ultrasound pressure (*F*_5,114_ = 0.37, *p* = 0.87). A two-way ANOVA revealed a significant effect of genotype× pressure interaction (both *p* < 10^−35^, two-way ANOVA). Although the response of Mec mutants appeared as if it might be modulated by pressure (Fig. 3B-D), this apparent modulation was not significant (*p* > 0.09, one-way ANOVA). Collectively, these results establish that behavioral responses to focused ultrasound depend on the TRN neurons and the MEC-4 protein.

Thus far, we have shown that focused ultrasound evokes reversal behaviors in freely moving *C. elegans* nematodes in a pressure- and stimulus duration-dependent manner (Fig. 2) and that such responses depend on the animal’s ability to detect mechanical, but not thermal stimuli (Fig. 3). Together, these findings imply that ultrasound exerts its effects on excitable tissues *via* a non-thermal, mechanical mechanism. Although additional work is needed to determine how ultrasound produces these effects, a leading possibility is the generation of mechanical strain in neurons expressing mechanosensitive ion channels like MEC-4.

MEC-4 is not the only protein thought to form a mechanosensitive ion channel in *C. elegans* nematodes. The TRP-4 protein is expressed in the CEP mechanoreceptor neurons and is an ortholog of the Drosophila NOMPC channel (Li et al., 2006) known to form mechanosensitive ion channels (Yan et al., 2013). A previous study showed that *C. elegans* responds to ultrasound-induced cavitation of microbubbles and proposed that these responses were due to action of TRP-4 (Ibsen et al., 2015). To determine if TRP-4 also contributed to ultrasound-evoked behaviors elicited in the absence of microbubbles, we analyzed the same *trp-4* strain used by Ibsen et al. (VC1141 *trp-4(ok1605)*). In agreement with the prior report (Ibsen et al., 2015), we observed a modest deficit in ultrasound-evoked behavior (Fig. 4A). A two-way ANOVA detected both a main effect of strain (*F*_1,228_ = 17.8, *p* < 0.0001) and a significant strain × pressure interaction (*F*_5,228_ = 4.8, *p* = 0.0003). The defect in these mutants was not specific for ultrasound-evoked behaviors, however: *trp-4* mutants had a lower average speed than wild-type mutants under baseline conditions (0.17 versus 0.21 mm/s, *p* = 0.0086, t-test, *n* = 20).

Because the *ok1605* allele encodes a partial in-frame deletion and because we also observed that these mutants grew slowly compared to wild-type animals, we tested two additional deletions in the *trp-4* gene: *gk341* and *sy695*. All three alleles, *ok1605*, *gk341*, and *sy695* are expected to encode deletions in the *trp-4* gene, which we verified by PCR analysis of genomic DNA (see Methods). Despite the expectation that that the three *trp-4* alleles would have the same ultrasound phenotype, we found that *gk341* and *sy695* mutants responded to ultrasound just like wild-type animals (Fig. 4B; two-way ANOVAs, main effects and interactions *p* > 0.29).

These findings suggested that the deficit in the VC1141 *trp-4(ok1605)* animals might be due to a mutation present in the genetic background. We tested this idea by out-crossing the *trp-4(ok1605)* animals against wild-type (N2) animals four times while tracking the *trp-4* mutation via PCR. Animals from this new strain, GN716 *trp-4(ok1605)*, had ultrasound-evoked behaviors that were indistinguishable from wild-type (Fig. 4B; two-way ANOVA, main effect and interaction *p* > 0.23). These results are summarized for the pressure of 1 MPa in Fig. 4C and suggest that the defect we and others (Ibsen et al., 2015) have observed in VC1141 *trp-4(ok1605)* animals is due to mutation/s in the genetic background of this strain.

**Figure 4.**
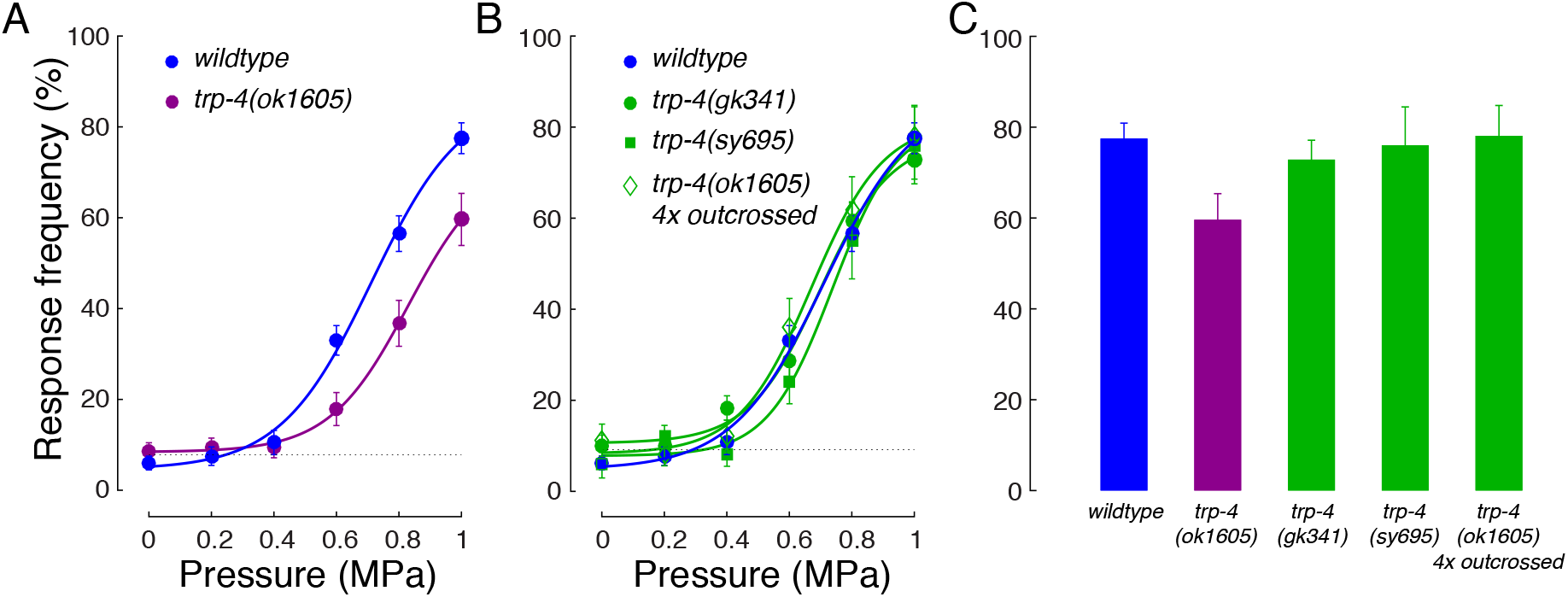
Strains carrying deletions in the *trp-4* NOMPC channel gene differ in their response to ultrasound stimulation. (A) Pressure-response curves of wildtype (blue) VC1141 *trp-4(ok1605)* (magenta) mutants used in a previous study (Ibsen et al., 2015). The smooth curve fit to the *trp-4(ok1605)* data yielded *F_max_* = 65%; *P1/2* = 0.83 MPa, slope = 0.13, base = 8%. (B) Pressure response curves of three other *trp-4* mutant lines were indistinguishable from wild-type. VC818 *trp-4(gk341)*, TQ296 *trp-4(sy695)*, and GN716 *trp-4*(*ok1605* mutants, which was derived from VC1141 by outcrossing four times with wild-type (N2) animals. (C) Response rate in four *trp-4* mutant strains. Bars are the mean (± s.e.m.) reversal rate evoked by ultrasound stimulation with the following parameters: 1kHz, 50% duty cycle, 200ms pulse duration, 1.0 MPa. Dotted line represents the average baseline response rate (see Materials and Methods). The number of animals analyzed across 10 trials is indicated in parentheses. We used PCR to verify that all strains harbored the expected deletions in the *trp-4* locus (Materials and Methods). Wildtype data are from Fig. 2C).

The finding that mechanosensation is an essential component of the biophysical effects of ultrasound suggests that there might be an optimal frequency of the ultrasound delivery that matches the mechanical properties of the tissue. We investigated this possibility by varying pulse repetition frequencies (PRFs) in the range from 30 Hz to 10 kHz, while keeping pulse duration, pressure, and duty cycle constant. We found that ultrasound indeed evoked reversals in a pulse repetition frequency-dependent manner (Fig. 5A). Response increased with PRF, reached a maximal value in the range of 300–1000 Hz, and decreased at higher frequencies. The shape of the curve is reminiscent of the prediction (Fig. 5A, green) of a model linking indentation to mechanical strain and MEC-4-dependent channel activation (Eastwood et al., 2015). We note that since stimuli were delivered at 50% duty cycle at all the tested frequencies, the same amount of energy was delivered at all pulse repetition frequencies. If the behavioral responses were the result of tissue heating, little or no modulation by the PRF would be expected. Yet, the plot shows and an ANOVA confirms a strong modulation of the response by the PRF (*F*_5,114_ = 10.8, *p* < 10^−7^).

We further hypothesized that discrete pulses may be more potent in eliciting mechanical effects because discrete pulses deliver multiple discrete mechanical events into the tissue. To test this idea, we varied the duty cycle while holding stimulus duration, pulse repetition frequency, and pressure values constant at 200 ms, 1kHz, and 1 MPa, respectively. Fig. 5B shows the relationship between response frequency and duty cycle. It reveals that a duty cycle of 50% was more than three-fold more potent than a continuous or 100% duty cycle protocol (77.5% compared to 24.0%, *p* < 10^−12^, t-test) even though a continuous stimulation delivers twice as much energy into the tissue as the pulsed protocol of 50% duty. In line with this finding, pulsed ultrasound stimulation has been found more effective than continuous stimulation in eliciting motor responses in rats (Kim et al., 2014). That study also found the value of 50% duty to be optimal. It is important to note that the 24.0% response rate for the continuous stimulus (100% duty) is above baseline (*p* < 0.001, t-test, *n* = 20). Thus, although a pulsed stimulus is more effective than a continuous stimulus, pulsing the ultrasound is not necessary to elicit a significant response. The figure further shows that the width of the individual mechanical events associated with the ultrasound can be quite brief—just 50 µs (5% of duty)—and still trigger appreciable behavioral responses (response rate of 34.0%, significantly different from baseline at *p* < 0.0001, t-test, *n* = 20). This is even though the energy delivered into the tissue is only 1/10th of that delivered at 50% duty.

**Figure 5.**
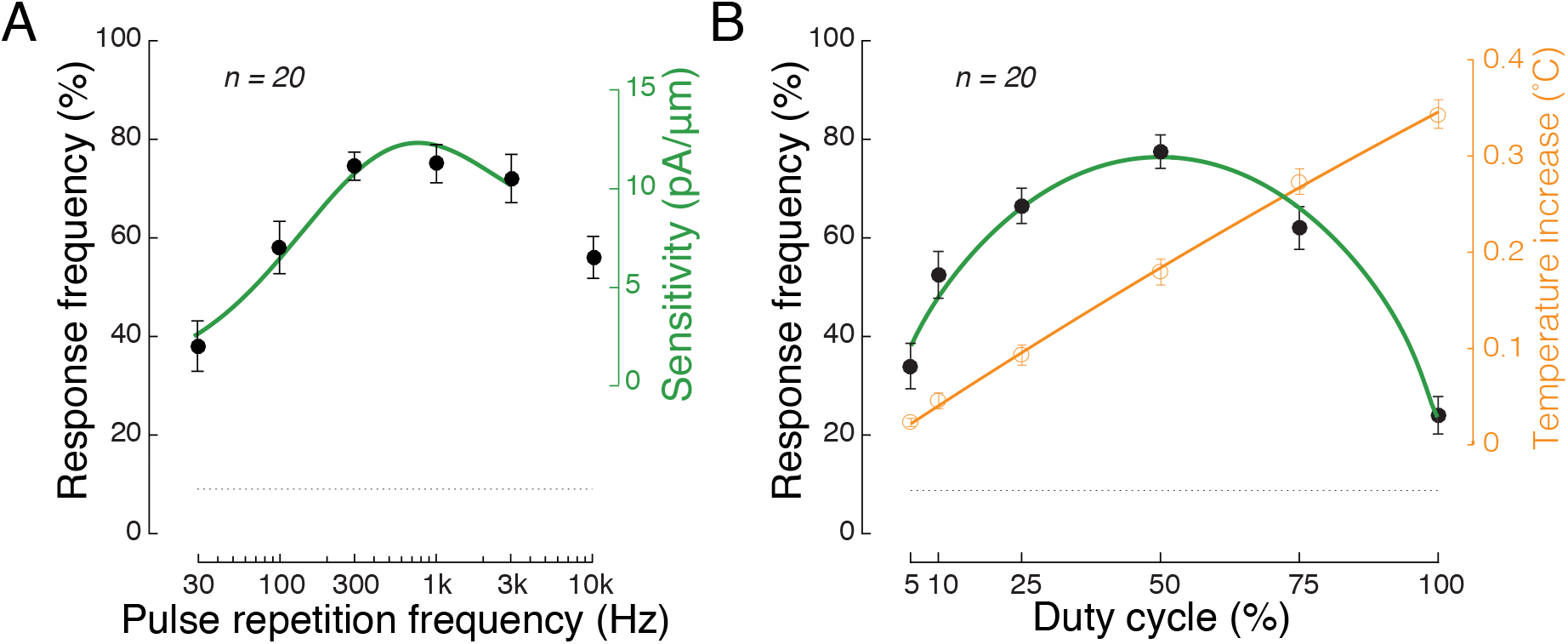
Ultrasound efficacy depends on pulse repetition frequency and duty cycle. (A) The response (black circles, mean±s.e.m.) of wild-type animals to a train of ultrasound pulses plotted as a function of pulse repetition frequency. The duty cycle was held constant at 50% in all cases, ensuring that all stimuli deliver the same amount of energy. The smooth curve (green) shows a simulation of the sensitivity of TRN currents to sinusoidal mechanical indentations (Eastwood et al., 2015). (B) The response (black circles, mean±s.e.m.) of wild-type animals as a function of duty cycle. Because the pulse repetition rate was 1 kHz in all cases, a duty cycle of 5, 10, 25, 50, 75, 100% corresponds to a pulse width of 50 µs, 100 µs, 250 µs, 500 µs, 750 µs, and 1 ms (continuous wave), respectively. The green curve shows the level of TRN activation expected from the frequency filtering shown in panel A (see Methods for details). Orange circles (mean±s.e.m., *n* = 5) show the increase in temperature observed as a function of duty cycle. The orange curve is a quadratic fit to the data included for visual clarity. For experiments in both A and B, the carrier frequency was 10 MHz, the stimulus amplitude was 1 MPa, and the stimulus duration 200 ms. The dotted line shows the baseline reversal frequency.

Whereas heating increases linearly with duty cycle (Fig. 5B orange plot), behavioral response frequency had a non-linear dependence on duty cycle. The heating effect is expected from the fact that the energy delivered in the tissue increases with duty cycle. To ask whether or not the dependence on duty cycle could be explained by the frequency dependence of TRN activation, we simulated the frequency distribution expected as a function of duty cycle and combined with this the model from Eastwood, *et al*. (Eastwood et al., 2015). This model matched the experimental results (Fig. 5B, green line). Collectively, the effect of duty cycle reinforces the idea that behavioral responses to ultrasound are mediated by mechanical effects and not by heating.

## Discussion

We sought to illuminate the biophysical mechanisms that underlie ultrasound stimulation of excitable cells. To do so, we used *C. elegans* as a model, harnessing its well-characterized and compact nervous system and comprehensive library of animals with specific genetic interventions. This animal has an extraordinary ability to detect tiny thermal fluctuations and mechanical stimuli (Ramot et al., 2008; O’Hagan et al., 2005). We found that ultrasound elicits robust reversal behaviors and that the response probability depends on stimulus intensity, duration, and specific pulsing protocols. Sensitivity to ultrasound and its modulation by pressure is preserved in mutants deficient in thermosensation and eliminated in mutants defective in mechanosensation. These findings are in agreement with a report (Zhou et al., 2017) suggesting that wild-type, but not *tax-4* mutants (which are defective in thermosensing (Ramot et al., 2008)) reverse in response to brief pulses of high-frequency ultrasound. Consistent with non-thermal, mechanical activation of sensory neurons linked to reversal behaviors, the response probability exhibited optima in both pulse repetition frequency and duty cycle.

Ultrasound-evoked reversal responses required expression of MEC-4, a key subunit of a touch-activated mechanosensitive ion channel. This finding implies that ultrasound can activate neurons by acting on mechanosensitive ion channels. Notably, because the MEC-4 ion channel complex is activated by mechanical forces and not by changes in membrane voltage or capacitance, our findings are inconsistent with the hypothesis that ultrasound acts by inducing changes in membrane capacitance (Krasovitski et al., 2011; Plaksin et al., 2014).

We tested whether other mechanosensitive channels might contribute to ultrasound-evoked behaviors by analyzing strains carrying deletions in the *trp-4* gene. Prior work showed that TRP-4 is a pore-forming subunit of mechanosensitive channel expressed by texture-sensing neurons in the worm’s head (Kang et al., 2010) and suggested that this protein could sensitize neurons to ultrasound stimulation. Yet, we did not detect any effect of the loss of TRP-4 channels on ultrasound-evoked responses in three independent strains carrying validated deletions in the *trp-4* gene. We did detect a decrease in ultrasound sensitivity in a fourth strain (VC1141) that was used in a previous study (Ibsen et al., 2015). However, this phenotype is not due to loss of *trp-4* function, since four rounds of outcrossing eliminated it. Rather, the partial loss of ultrasound sensitivity is likely to be due to unidentified mutation(s) in the VC1141 strain. Additional investigations will be needed to identify the affected gene(s), an effort that could reveal additional genetic factors regulating sensitivity to ultrasound stimulation.

This study provides evidence that ultrasound can stimulate neurons through its mechanical mode of action. Within the mechanical domain, there can be several specific candidate mechanisms at play. First, ultrasound may elicit cavitation, a phenomenon characterized by formation and collapse of gaseous bodies in liquid media or soft tissues. However, for frequencies above 1 MHz, cavitation requires pressures greater than 5 MPa and the cavitation threshold for 10 MHz is even higher (Nightingale et al., 2015). Thus, both the 10MHz transducer and low pressures we used make cavitation unlikely. Second, the incident tissue, such as a cell membrane, experiences oscillations with period equal to the ultrasound carrier frequency.

The pressures used for neuromodulation can cause appreciable particle displacement (on the order of 0.01– 0.1 *µ*m (Gavrilov et al., 1976)). Nonetheless, the displacement is distributed in sinusoidal fashion along the wavelength (about 100 *µ*m at 10 MHz) of the propagating wave. This creates a very small displacement gradient (e.g., 0.1 *µ*m per 100 *µ*m). It is questionable whether such a small gradient can cause significant enough deformation of a pore segment of an ion channel with regard to the channel dimensions. Moreover, the primary pressure oscillations, which occur at a specific carrier frequency, cannot explain the frequency dependence of the responses (Fig. 5A). The third and most probable form of mechanical energy underlying the effects in this study is the acoustic radiation force (Trahey et al., 2004; Sarvazyan et al., 2010; Iversen et al., 2017). Acoustic radiation forces result from differences in acoustic intensities at individual points in space. The differences can be caused by ultrasound absorption, scattering, reflection, or other phenomena, and lead to net forces on the tissue (Duck et al., 1998). Acoustic radiation force exerts a steady pressure on a target throughout the time of ultrasound application. This steady pressure may stretch a cell membrane to an extent that affects conformation states of ion channels or other active molecules tied to the membrane. A simulation of the propagating ultrasound field, for the pressure of 1 MPa, revealed that a net acoustic radiation force of 873 nN can act on the animal‘s head (see Materials and Methods). This exceeds the animal’s sensitivity threshold to mechanical forces, which is 50-100 nN (Petzold et al., 2013). Thus, the acoustic radiation force expected for a 1 MPa stimulus is sufficient to engage the animal’s mechanosensation. It is worth to stress that the radiation force acts during the On epochs of the ultrasound (black rectangles in Fig. 1C), and not during the Off epochs when the ultrasound amplitude is zero. This way, pulsed ultrasound delivers force pulses at a specific pulse repetition frequency, and there can therefore be a modulation by the pulse repetition frequency (Fig. 5A).

Our findings implicate mechanical force as a major physical effect of ultrasound on neurons and their ion channels, and delineate a complete pathway from mechanical force to activation of excitable cells. In this light, it is tempting to reiterate a potential unifying mechanism linking ultrasound to activation of excitable cells (Tyler, 2011). Suppose that ultrasound exerts similar mechanical effects in complex nervous tissues as it does in *C. elegans*. In addition, suppose that ultrasound deforms tissue and generates mechanical strain in neurons sufficient to activate mechanosensitive ion channels, as in *C. elegans*. Ion channels likely to subserve this function in mammals include the intrinsically mechanosensitive K2P family of potassium channels (Brohawn et al., 2014) and piezo channels (Syeda et al., 2016) known to be expressed in the brain. In support of this idea, the activity of K2P channels, including TREK-1, TREK-2, and TRAAK, is potentiated by ultrasound stimulation in heterologous cells (Kubanek et al., 2016). Sensitivity to the mechanical effects of ultrasound might not be limited to primarily mechanosensitive channels. For instance, voltage-gated sodium channels have been implicated in activation of neurons by ultrasound (Tyler, 2011; Tyler, 2012; Kubanek et al., 2016) and are known to be sensitive to membrane tension (Beyder et al., 2010).

Exactly how ultrasound stimulation is translated into local mechanical strain will depend on the material properties of the excitable tissues under study. In the case of *C. elegans*, we found that the pulse repetition frequency and duty cycle dependence of ultrasound-evoked behaviors agreed (Fig. 5A) with a model that links tissue indentation associated with a mechanical stimulus to the activation of MEC-4-dependent channels (Eastwood et al., 2015). The correspondence (Fig. 5A) suggests that ultrasound-induced radiation force elicits a profile of mechanical strain similar to that produced by indentation with a physical probe. The model (Eastwood et al., 2015) also provides insight into why stimuli of 50% duty are the most potent (Fig. 5B): stimuli delivered at lower or higher duty cycle values contain a majority of their energy in high-frequency harmonics (relative to the 1 kHz pulse repetition frequency), and these high-frequency harmonics are filtered out by the tissue (Fig. 5A). However, the relationship between bursts of ultrasound pulses and the mechanical strain we infer it generates is not currently known and future measurements of ultrasound-induced strain will be needed to fill this knowledge gap for worms and other tissues. Consistent with the proposal that tissues differ in their mechanical filtering properties, ultrasound-evoked behaviors have an optimal pulse repetition frequency near 500 Hz in *C. elegans* and show strong effects at about 3000 Hz (King et al., 2013) in mice. These results indicate that an improved understanding of mechanical filtering by soft tissues will be needed to further the long-term goal of applying ultrasound as a noninvasive modality to stimulate excitable cells.

This study reveals that the pulsatile forces associated with ultrasound are potent enough to activate mechanically sensitive ion channels in living animals. Given that many ion channels expressed in neurons and glia are mechanically sensitive (Ostrow and Sachs, 2005; Tyler, 2012), this study illuminates one way that ultrasound could influence activity in the brain. Another way would be to sensitize specific neurons to ultrasound through ectopic expression of a channel known to be mechanosensitive, a strategy referred to as sonogenetics (Ibsen et al., 2015). In both scenarios, our work underscores the importance of tuning stimulus parameters to maximize acoustic radiation force and may help to further the development of ultrasound as a non-invasive and spatially precise tool to study the nervous system and potential to ameliorate neurological disorders.

## Acknowledgements

We thank our colleagues at Stanford, Z. Liao, M. Menz, P. Ye, M. Prieto for laboratory and technical assistance and input on the experimental design; and M. Maduke, P. Khuri-Yakub, K. Butts-Pauly, M. Menz, M. Prieto, and J. Brown for helpful comments. We also thank Z. Pincus (Washington University, St. Louis) for providing animals for initial pilot experiments. A. Sanzeni and M. Vergassola for additional model calculations. This work was supported by NIH grants K99NS100986 (JK), R01NS047715 and R01NS092099 (MBG), gifts from the Stanford Neuroscience Institute and Mathers Foundation (to MBG, SB) and a Stanford Medicine Dean’s fellowship (to JK).

